# Metabolomics of Mouse Embryonic CSF Following Maternal Immune Activation

**DOI:** 10.1101/2023.12.06.570507

**Authors:** Boryana Petrova, Tiara E Lacey, Andrew J Culhane, Jin Cui, Alexander Raskin, Aditya Misra, Maria K Lehtinen, Naama Kanarek

## Abstract

The cerebrospinal fluid (CSF) serves various roles in the developing central nervous system (CNS), from neurogenesis to lifelong cognitive functions. Changes in CSF composition due to inflammation can impact brain function. We recently identified an abnormal cytokine signature in embryonic CSF (eCSF) following maternal immune activation (MIA), a mouse model of autism spectrum disorder (ASD). We hypothesized that MIA leads to other alterations in eCSF composition and employed untargeted metabolomics to profile changes in the eCSF metabolome in mice after inducing MIA with polyI:C. We report these data here as a resource, include a comprehensive MS^1^ and MS^2^ reference dataset, and present additional datasets comparing two mouse strains (CD-1 and C57Bl/6) and two developmental time points (E12.5 and E14.5). Targeted metabolomics further validated changes upon MIA. We show a significant elevation of glucocorticoids and kynurenine pathway related metabolites. Both pathways are relevant for suppressing inflammation or could be informative as disease biomarkers. Our resource should inform future mechanistic studies regarding the etiology of MIA neuropathology and roles and contributions of eCSF metabolites to brain development.

## Introduction

The cerebrospinal fluid (CSF) provides a protective fluid cushion for the brain (Fame & Lehtinen, 2020; Lacey *et al*, 2023)and delivers neuroactive proteins, peptides, and small molecules that are critical for normal brain development (Saunders *et al*, 2023; Gelb & Lehtinen, 2023). In the case of the developing cerebral cortex, whose neural progenitor cells line the brain ventricles and extend primary cilia into the CSF, age-appropriate cocktails of CSF growth factors help ensure that progenitors remain healthy, acquire the correct identity, and proliferate in a developmentally appropriate manner(Chau *et al*, 2018; Lehtinen *et al*, 2011; Kim *et al*, 2010). Conversely, the CSF can be used as a readout of different stages of brain development including metabolic state (Chau *et al*, 2018; Fame *et al*, 2019). Indeed, CSF is commonly sampled for biomarkers of infections and neurologic diseases.

While much of the CSF is produced by the choroid plexus (ChP), an epithelium located in each brain ventricle (Gelb & Lehtinen, 2023; Saunders *et al*, 2023), substances can seep into the CSF from the brain’s interstitial fluid or any of the cells lining the ventricles. Given the importance of CSF composition for brain development, new techniques and mouse models have recently been harnessed to elucidate the transcriptome, proteome, and some diagnostic biomarkers in the CSF during development (Saunders *et al*, 2023). While meaningful efforts have characterized some small molecules in the CSF as critical for CSF function, i.e., nutrients, salts, and signaling molecules, the CSF metabolome remains relatively less characterized.

To date, efforts to establish a reference (normative) adult mouse CSF metabolome (Wishart *et al*, 2008; Stoop *et al*, 2010; Mandal *et al*, 2012) have identified a growing list of CSF metabolites associated with adult neuroinflammation as well as neurologic and psychiatric phenotypes (Panyard *et al*, 2021). However, the CSF metabolome during development has yet to be comprehensively described. This gap in knowledge compromises our understanding of normal development as well as the etiology of neurodevelopmental disorders.

Here we describe the profiling of the embryonic CSF (eCSF) metabolome and share this resource with the community of developmental biologists and neuroscientists. Utilizing liquid chromatography-mass spectrometry (LC-MS), we generated a comprehensive database of embryonic day (E)14.5 mouse eCSF of CD-1 mice, including mass verification at the level of mass-to-charge (MS^1^) or at the level of fragmentation spectrum (MS^2^). This database can be used as a reference library for future studies, and indeed we utilized it to study healthy mice from two strains (C57BL/6 and CD-1), two developmental time points (E12.5 and E14.5), and embryos that were exposed to maternal immune activation (MIA) by polyinosinic-polycytidylic acid (polyI:C, a synthetic viral mimetic) injection of pregnant dams. We chose this model of inflammation-induced neuropathology because a growing body of evidence suggests that changes in eCSF composition following maternal illness are likely to contribute to negative sequelae in offspring (Parada *et al*, 2005, 2006; Panyard *et al*, 2021; Muk *et al*, 2022; Chau *et al*, 2015), such as neurocognitive impairment, schizophrenia, and autism spectrum disorders (ASD) (Ashwood *et al*, 2006; Patterson, 2009; Knuesel *et al*, 2014; Estes & McAllister, 2016; Lu-Culligan & Iwasaki, 2020).

The choice of time points and mouse strains were led by relevance to the MIA mouse model: The E12.5 developmental timepoint coincides with the peak levels of pro-inflammatory cytokines in maternal plasma 3 hrs following MIA induction by exposure to polyI:C. The E14.5 developmental timepoint coincides with reported abnormalities following MIA induction of brain cytokine levels including IL-6 (Smith *et al*, 2007; Wu *et al*, 2017; Garay *et al*, 2013; Hsiao & Patterson, 2011) and IL-17a (Hsiao *et al*, 2012; Kim *et al*, 2017; Choi *et al*, 2016). Additionally, we previously reported a proinflammatory eCSF state at E14.5 that includes elevated levels of CCL2 in the eCSF that is accompanied by the accumulation of immune cells (Panyard *et al*, 2021; Cui *et al*, 2020).

We observed differential abundance of metabolites from two metabolic pathways related to inflammation: glucocorticoids (Gaber *et al*, 2017) and the kynurenine pathway (Murakami *et al*, 2021, 2023; Cervenka *et al*, 2017). Glucocorticoids are anti-inflammatory and immunosuppressive, and among the most prescribed drugs for the treatment of immune and inflammatory disorders (Cruz-Topete & Cidlowski, 2014). Yet, glucocorticoids (i.e., corticosterone, the main stress hormone utilized in rodents) can be either pro- or anti-inflammatory at different developmental stages (Brown *et al*, 1996; Wyrwoll *et al*, 2011; Astiz *et al*, 2020). The function of glucocorticoids in the eCSF, and potential role(s) of these small molecules in shaping neurocognitive effects of maternal inflammation are not known.

Another inflammation regulatory metabolic pathway is the kynurenine pathway. Kynurenine is a downstream metabolite in the catabolism of the essential amino acid tryptophan. Tryptophan is a precursor to serotonin (5-HT), a neurotransmitter with key roles in regulating neurodevelopmental processes including circuit formation (Bonnin *et al*, 2007, 2011; Goeden *et al*, 2016). However, depending on the tissue or context, it is estimated that up to 90% of tryptophan is catabolized to kynurenine (Dehhaghi *et al*, 2019), a well-documented anti-inflammatory metabolite with direct effects on both brain and immune function (Dantzer, 2016; Davis & Liu, 2015; Cuzzell, 1986). Other metabolites in the kynurenine metabolism pathway, e.g. kynurenic acid and quinolinic acid, are also immunomodulatory (Savitz, 2020). Kynurenine, kynurenic acid, and quinolinic acid are all altered across the maternal-fetal axis (maternal liver, embryonic serum, and brain) following maternal immune activation (Goeden *et al*, 2016). Notably, altered kynurenine pathway metabolism in the adult has been associated with schizophrenia (Erhardt *et al*, 2001; Schwarcz *et al*, 2001; Sathyasaikumar *et al*, 2011; Linderholm *et al*, 2012) and bipolar disorder with a history of psychosis (Olsson *et al*, 2012).

Our eCSF metabolomics database provides an important resource for guiding investigations of neural development. Additionally, we uncovered several metabolic aberrations induced by MIA, including increased levels of metabolites of the kynurenine pathway and rodent corticoid stress response across the maternal-fetal axis. Our work may reveal targetable pathways for eventually developing CSF-based interventions for at-risk offspring following maternal illness.

## Results

### Generation of an embryonic CSF metabolome database

We applied untargeted metabolomics to generate a metabolite library of polar metabolites in mouse eCSF at embryonic day (E)14.5. eCSF metabolomics is challenging because it must be accomplished with very small sample volumes, and analysis must account for the low signal-to-noise ratio that stems from the relatively low metabolite concentrations in the eCSF compared to other biofluids. We therefore optimized a consistent, reliable sample collection and analysis approach: In LC-MS analysis, a signal, also called a feature, is defined as an m/z value with an associated retention time. Due to the high sensitivity of mass spectrometry, background and contaminant signals with defined m/z’s and retention times can sometimes be indistinguishable from true biological signals. This noise becomes especially confounding when a sample like CSF contains a low abundance of metabolites. Therefore, a rigorous analytical method is required to ensure that our metabolite library construction is based on true biological signals from eCSF.

To that end, we compared the ability of two analytical approaches to identify non-biological background (noise), namely 1) mock filtering and 2) “IROA” (Isotopic Ratio Outlier Analysis). Our goal was to filter out this noise from our analyses of the eCSF metabolome (Fig 1A and B). In the “mock filtering” approach, we compared the eCSF profile to multiple mock samples. The mock samples were empty tubes that underwent the exact same series of sample preparation steps through injection into the LC-MS, and thus they only differed in the actual presence of eCSF in the collection tubes. This approach exposed background features that were shared by the mock samples and eCSF samples, including technical noise and signals of non-biological contaminants. These signals were excluded from our defined eCSF metabolome. True eCSF-specific signals were defined as signals with 3-fold higher abundance in eCSF compared to mock samples (Fig. 1C). In the “IROA” approach (Qiu *et al*, 2018), we employed a ^13^C-labeled reference metabolome to credential detected signals (Fig EV1). Briefly, in this approach, we first mixed the eCSF metabolome with a ^13^C-labeled reference metabolome, a metabolome where most naturally occurring ^12^C carbon atoms have been substituted with ^13^C, a heavier stable isotope of carbon. Because LC-MS resolution allows detection of the common ^12^C-metabolites side-by-side with the reference ^13^C metabolites, detected signals in the eCSF could be credentialed as “true biological signals” only if they had matched a heavy signal in the reference metabolome. All other signals were excluded from the analysis (Fig 1D). Only ionized signals are detected by the MS; these can form either by deprotonation (negative mode), or by the addition of a proton (positive mode). Some metabolites tend to be better detected in one mode vs the another, and in the case of small molecules, they tend to acquire a single charge. To capture the maximum number of metabolites, we analyzed both positive and negative modes. Both the mock filtering and IROA approaches excluded a similar order of magnitude number of signals: mock filtering retained 16% (negative mode) and 18% (positive mode) of total features remained after mock filtering (Fig 1C) while IROA-based filtering retained 12% (negative mode) and 16% (positive mode) (Fig 1D). Overall, we detected more signals using the mock filtering approach with 632 (negative mode) and 798 (positive mode) features being detected using the mock-filtered compared to the 35 (negative mode) and 71 (positive mode) detected using IROA. We concluded that the IROA approach is too stringent for eCSF metabolite profiling, likely due to the use of a yeast metabolome IROA library rather than a CSF-specific labeled metabolome, which is not currently available. Therefore, we proceeded to apply the mock filtering approach for all subsequent analyses.

**Figure 1.**
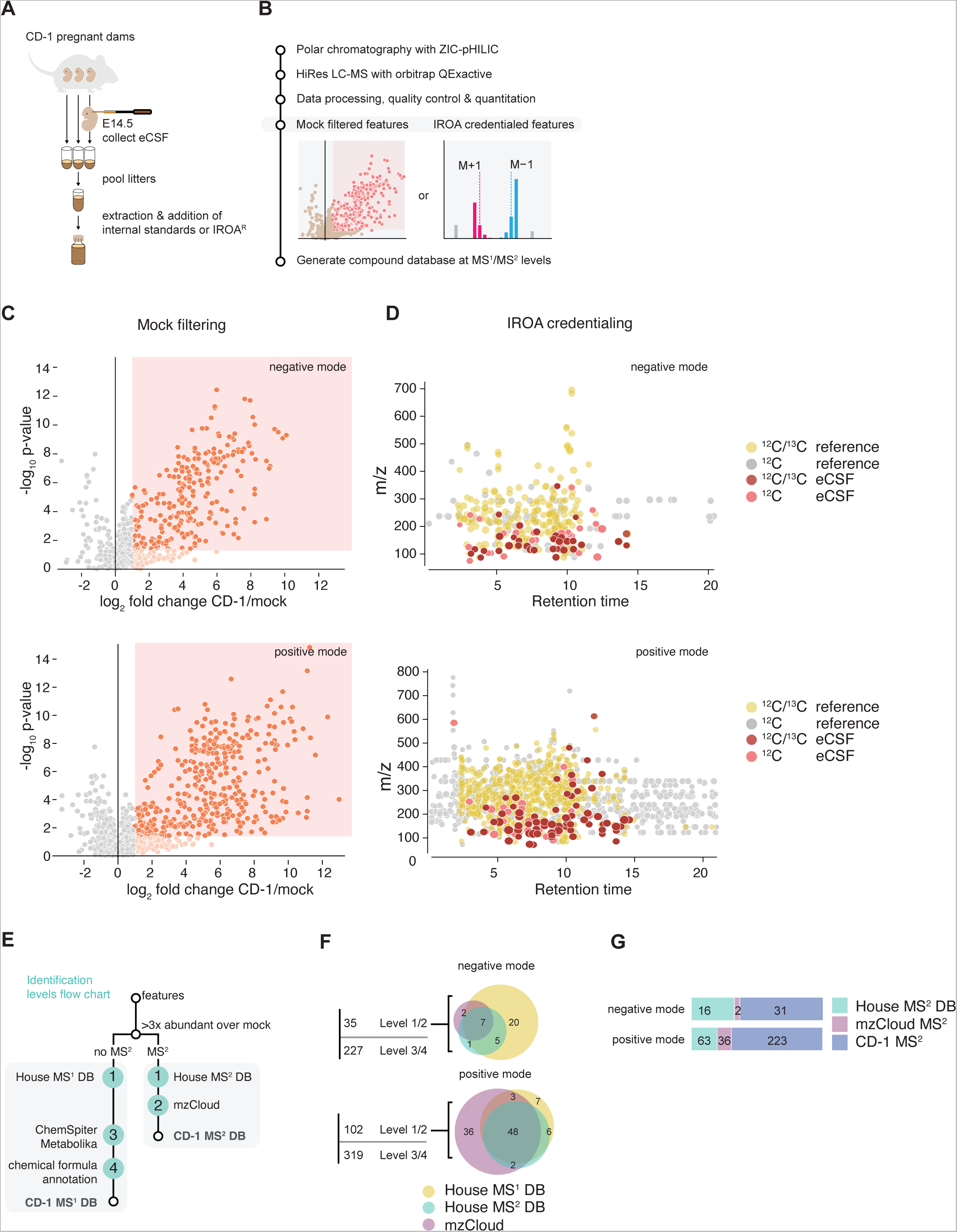
An approach to generate an embryonic CSF (eCSF) compound database as a tool for future developmental and neurological research. A Schematic depicting experimental workflow for embryonic CSF sample preparation. CSF from embryos (eCSF) from a single pregnant mouse was pooled and 3-7 µl extracted for analysis. Each pregnant mouse was considered a replicate. B Schematic depicting the analytical workflow in generating an eCSF polar compound database. Polar chromatography was performed in HILIC mode (ZIC-HILIC: zwitterionic hydrophilic interaction column) and metabolites were detected using a high-resolution (HiRes) mass spectrometer (Thermo Orbitrap QEactive). Data was processed using two separate approaches depending on compound identification strategy. Contaminants were filtered either based on an extensive set of mock samples or on labeled metabolome correspondence (“IROA” credentialing, see also Fig EV1). Signals (features) passing filtration and quality control were then used to generate compound databases at MS^1^ and MS^2^ level. C Representative results from untargeted polar metabolomics on eCSF from CD-1 mice (N=7) for mock-filtered dataset outlined in B. Positive- and negative-mode analysis is shown separately. Raw data for polar metabolomics was processed using CompoundDiscoverer (CD) 3.3. Mock and eCSF samples are compared using a volcano plot. Signal was normalized based on targeted analysis of polar metabolites and mean-centering per sample (see methods for details). Metabolites highlighted in orange are >2-fold significantly higher in eCSF than in mock samples. D Representative results from untargeted polar metabolomics on eCSF from CD-1 mice (N=7) for IROA credentialed dataset as outlined in B. Relevant biological features were identified using IROA-“LTRS” – a 1/1 mix of 95:5/5:95 unlabeled (^12^C) and ^13^C-labeled reference yeast metabolome (^12^C/^13^C reference, IROA® TruQuant Yeast Extract Semi-targeted QC Workflow). IROA-credentialed features were identified from eCSF (^12^C, unlabeled) mixed with a reference internal standard (“IS”) that was a mix of 5:95 unlabeled (^12^C) and ^13^C-labeled metabolome. Non-credentialed ^12^C signals (^12^C reference or ^12^C eCSF), with no matching IROA ^13^C signals, are depicted as indicated on legend. E Strategy for the implementation of the eCSF library into an untargeted analysis workflow combining in-house and online databases at MS^1^ and MS^2^ levels. Levels of identification certainty (1 through 4) are depicted following recommendations from the metabolomics community. G Breakdown of the composition of the eCSF-specific filtered features from the mock-filtered CD-1 untargeted dataset for which we have obtained MS^2^ level information. Negative and positive mode are presented independently. Matches to in-house (House MS^2^ DB) and external (mzCloud^TM^ MS^2^) libraries are depicted. The remaining spectra were manually quality controlled and consolidated into the CD-1 MS^2^ database.

For careful annotation and identification of detected metabolites, we employed a scoring system based on community recommendations for best practices in reporting untargeted metabolomics datasets (Sumner *et al*, 2007; Members *et al*, 2007; Salek *et al*, 2013) (Fig 1E). We applied the scoring system on features detected in eCSF in both negative and positive modes (Fig 1F). We incorporated into our workflow extensive in-house standard libraries reference, both at MS^1^ and MS^2^ levels, as well as information from several online databases. Level 1 reflects the highest level of certainty, and it integrates chemical information from retention time matching with MS^1^ (high resolution exact mass) and/or MS^2^ data from in-house standards reference libraries that are chromatographic method-specific libraries. At this level of identification, if a detected feature matched a standard metabolite from our MS^1^ in-house libraries, we denoted it as a “House_MS^1^_DB”, whereas if it matched MS^2^ in-house libraries, we denoted it as “House_MS^2^_DB” (Fig 1F). For Level 2 annotation, we relied on the mzCloud^TM^ online spectral database as our reference, and because no specific chromatographic method retention times are available when using mzCloud^TM^, this level reflects lower certainty in compound identification. Yet, utilization of mzCloud^TM^ extended our compound annotation and added 2 metabolites in negative mode and 36 metabolites in positive mode to our defined eCSF metabolome (Fig 1F). Finally, all remaining compounds for which we were able to obtain a chemical formula (227 in the negative mode and 319 in the positive mode) were annotated at certainty Level 3 and 4 and given molecular weight (MW) and retention time (RT) labels. In cases when a chemical formula matched entries from external compound databases (Metabolika or ChemSpider), the name of the top listed hit from the database was used and the level of certainty was depicted as Level 3 (putative annotation). For many metabolites that were scored as Level 3, we also obtained MS^2^ data (31 of 49 in negative mode and 223 of 322 in positive mode, Fig 1G). This could be extremely helpful with metabolite identification in future studies using our spectral database. All detected signals with insufficient intensity to assign a molecular formula were excluded from our database. Although we could not verify the identity of all annotated signals (below Level 1), our data suggest that these are eCSF-specific, relevant metabolites and not contaminants. However, we cannot exclude the possibility that some are artifacts from oxidation, chromatography, and ionization, these signals could be degradation products or other chemical derivatives of other compounds. Therefore, care must be taken in interpreting results and extensive follow up experiments will be required to fully annotate the eCSF-specific metabolome. In sum, as resource for future studies, we report here a stringent list of metabolites detected in the mouse eCSF at day E14.5, with assigned annotation, and transparent report of annotation confidence.

### Comparative untargeted eCSF metabolomics in two mouse strains and at two developmental time points

Brain and liver tissue from different mouse strains can exhibit distinct baseline metabolomic patterns (Burlikowska *et al*, 2020). This is relevant to the MIA model because behavioral and immunological effects are also strain-dependent (Schwartzer *et al*, 2013; Kentner *et al*, 2019). To broaden the applicability of our eCSF metabolomics database from CD-1 mice, we expanded our untargeted metabolite profiling to also include C57Bl/6 mice. MIA has been traditionally modeled in mice with a C57BL/6 background due to the requirement for the presence of commensal segmented filamentous bacteria (SFB) in the maternal gut microbiome for the induction of the MIA phenotypes in offspring (Kim *et al*, 2017). We have shown that SFB is also present in the maternal gut microbiome of pregnant CD-1 mice (Cui *et al*, 2020). In addition, we have shown in CD-1 mice that MIA induction by injection of the synthetic viral genome mimetic polyI:C at embryonic day E12.5 leads to a pro-inflammatory eCSF cytokine profile and infiltration of immune cells through the embryonic ChP 48 hours later, at E14.5 (Cui *et al*, 2020). This study and others illustrate the relevance of both C57BL/6 and CD-1 mouse strains for the field of MIA research. Therefore, we profiled both these mouse strains to provide extensive reference data for future studies of eCSF of both CD-1 and C57BL/6 mouse strains.

We queried signals present in both CD-1 and C57Bl/6 eCSF samples using our in-house compound databases and the CD-1-eCSF database we curated (CD-1-DB, presented in Fig 1) at MS^1^ and MS^2^ levels. We matched features at both MS^1^ and MS^2^ levels with two reference libraries: our “CD-1” and “in-house” libraries. More features matched to CD-1 library than to the in-house library and there was a good overlap between the two (Table 1 and Fig 2A).

**Figure 2.**
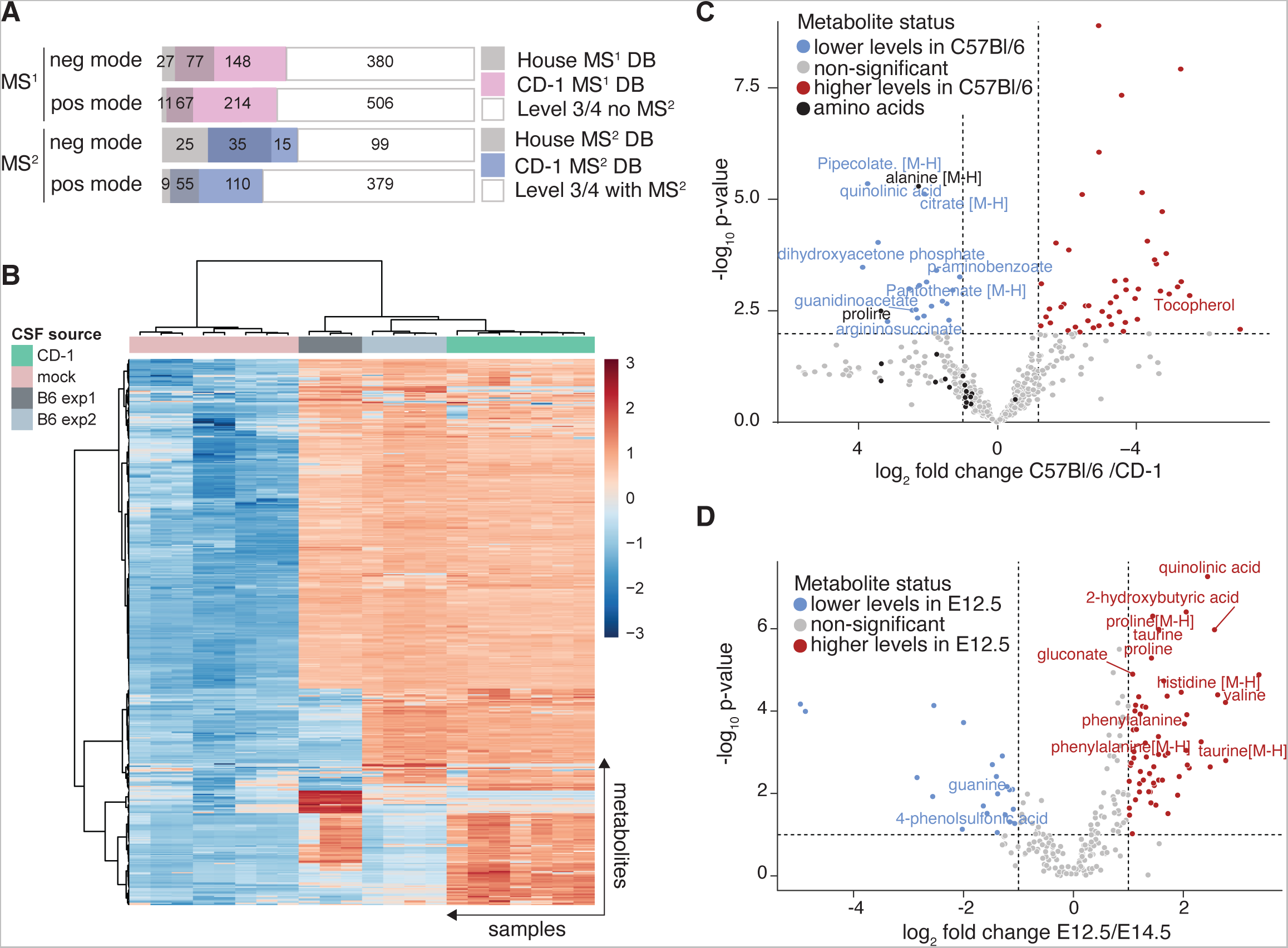
Application of eCSF library to maternal immune activation (MIA) model. A Breakdown of the composition of the eCSF-specific filtered features based on an overlap between CD-1 and C57BL/6 eCSF untargeted metabolome at the corresponding embryonic stage (E14.5). Two independent C57BL/6 eCSF cohorts from two mouse strains were treated as distinct batches because sample collection was performed on different days. Filtered untargeted features from C57BL/6 with either an in-house database match (House MS^1^ DB) or a CD-1 database match (CD-1 MS^1^ DB) are shown independently for MS1 or MS2 databases as indicated. Remaining features, with or without MS^2^-level information, which were not present in our databases, were identified at Level 3 and 4 and were also indicated. B Heatmap for the combined positive and negative mode data in A. Mock samples were included for comparison. Further 5% (Interquantile range) filtering step, log-transformation and Pareto scaling were performed within the MetaboAnalyst platform. C Volcano plot for the combined positive and negative mode data in A. For clarity, only significantly changed metabolites between CD-1 and C57BL/6 eCSF with in-house library match were annotated and also only negative ion adducts are indicated. In addition, all detected amino acids are indicated, with only significantly altered (alanine and proline) annotated. D Volcano plot, depicting a comparison between polar metabolites detected by untargeted metabolomics from eCSF (C57BL/6 samples) of E12.5 and E14.5. Compounds are Level 1 to 3 were used to generate the plot. For clarity, only significantly changed metabolites with in-house library match were annotated and also only negative ion adducts are indicated. E Schematic depicting the collection strategy of maternal serum for cytokine quantitation by ELISA upon injection of polyI:C and induction of MIA.

**Table 1.**
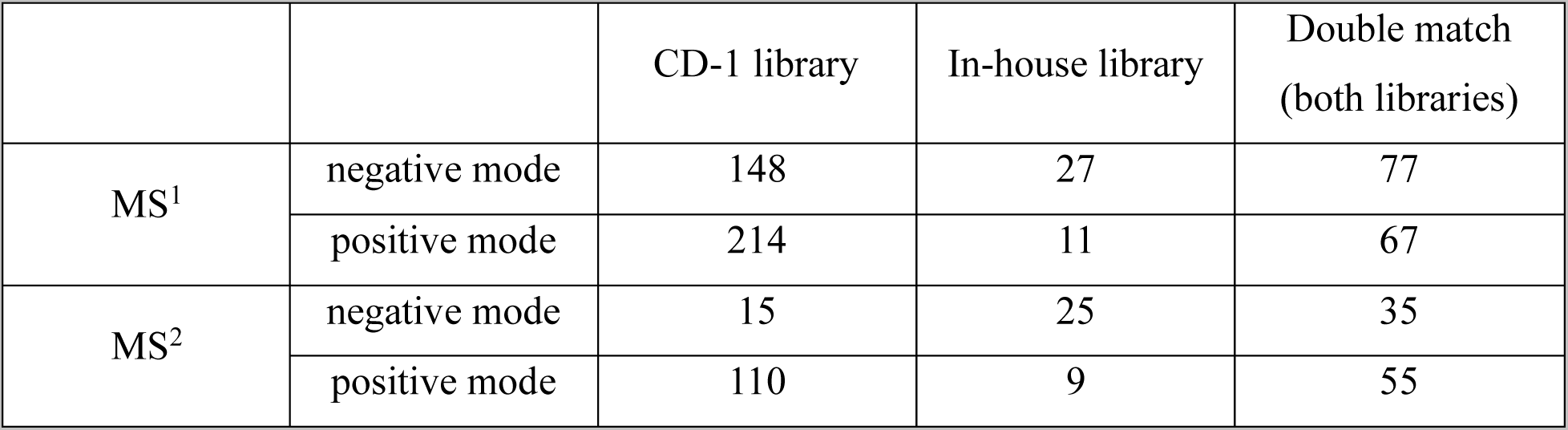
Number of features from CD-1 and C57Bl/6 eCSF samples identified as matching features to the reference metabolite libraries “CD-1” and “in-house”.

|  |  | CD-1 library | In-house library | Double match<br>(both libraries) |
| --- | --- | --- | --- | --- |
| MS<sup>1</sup> | negative mode | 148 | 27 | 77 |
|  | positive mode | 214 | 11 | 67 |
| MS<sup>2</sup> | negative mode | 15 | 25 | 35 |
|  | positive mode | 110 | 9 | 55 |

To address the question of overlap between the two different mouse strain, we compared the total number of filtered features to total features in the CD-1-DB. We detected 270 negative mode metabolites in the C57Bl/6 eCSF metabolome, out of the 421 metabolites present in our negative mode CD-1-DB (64%). Similarly, we detected 198 positive mode metabolites in the C57Bl/6 eCSF metabolome, out of the 262 metabolites present in the positive mode CD-1-DB (76%). We assumed that at least some of the metabolites that were detected in eCSF from CD-1 mice but not in C57Bl/6 mice could be missing from the C57Bl/6 samples for technical, and not biological reasons (e.g. LC-MS related retention time shifts or batch effect dependent changes in sensitivity of detection). We applied a highly stringent filter in our analysis that is based on comparison of the samples to the integrated sum of false signals from an extensive number of mock samples. We applied this stringent filter because we prioritized reduction of false positive results over false negative results. Therefore, this step likely resulted in exclusion of metabolites that varied between the two mouse strains likely due to batch effects. Nevertheless, biological reasons could dictate dissimilar gene expression and metabolism both in the mothers and in the embryos in these two strains. The hundreds of metabolites that were detected in both mouse strains allowed us to investigate in more detail the similarities and differences between the CD-1 and C57Bl/6 eCSF metabolomes (Fig 2A and B, Fig EV2A and B). Some metabolites (such as the abundant amino acids) were not significantly different between the two mouse strains, while others were (Fig 2C). Among the significantly different metabolites were Level 3 and 4 putatively annotated compounds that were present in our CD-1-DB but not in our in-house or online databases. These metabolites that we annotated and scored in our eCSF database represent potentially novel compounds that could introduce interesting metabolic differences to be considered in future eCSF research.

Next, we focused on C57Bl/6 mice and applied our eCSF database to assist untargeted metabolomics characterization of eCSF from two embryonic stages relevant for the MIA model, E12.5 and E14.5 in (Fig 2D). Collection of eCSF from these two developmental stages was performed and run on our instruments on different days, which may cause experimental variability due to batch effects. Therefore, to minimize technical and experimental variability, we combined data from two independent experiments (each with eCSF collected from a minimum of three independent litters). We detected differential abundance of several metabolites between E12.5 and E14.5 (Fig 2D), including the amino acids taurine, histidine, proline, phenylalanine, and valine. These amino acids were abundantly detected, and their differential levels reflect metabolic changes that occur over the course of brain development. Amino acids that were reported to be significantly altered in the fetal brain across development and during hyperglycemia in maternal diabetes, such as aspartate, asparagine, glutamate, and GABA (Perez-Ramirez *et al*, 2023), were not different in the eCSF in E12.5 vs. E14.5. These results reflect the distinct metabolic compositions of brain tissue and the CSF, underscoring the unique metabolic composition of the eCSF that we explored here and the importance of including the unique CSF metabolome rather than relying on brain parenchymal changes to infer CSF values. The dataset presented here can further serve the community as a reference for eCSF specific metabolites and inform on metabolic alterations during this critical stage of embryonic development.

### eCSF metabolomics following MIA reveals unique metabolite composition

To comprehensively profile metabolic changes in eCSF following induction of inflammation in the mother, we collected eCSF 3h and 48h following polyI:C delivery (Fig 3A, F). We confirmed elevated cytokine levels in polyI:C exposed dams at 3h (Fig EV3A and B) that recovered to near baseline levels by 48h following polyI:C exposure (Fig EV3C and D). We independently processed eCSF samples from two independent experiments per time point and analyzed them by untargeted metabolomics. We detected significant changes among eCSF polar metabolites between embryos collected from saline-versus polyI:C-injected dams at both time points (Fig EV3E and F). We observed biological variability between our independent experiments, as has been reported for the maternal response to polyI:C previously (Meyer *et al*, 2006; Kowash *et al*, 2019). At 3h following polyI:C delivery, we noted a consistent change in a cortisol-related metabolites (Fig 3B-E). In addition, at 3h and at 48h we observed several changes in the kynurenine pathway (Fig 3B-J) including the metabolites indoxyl sulfate, kynurenine, quinolinic acid and niacinamide. The changes in metabolites of the kynurenine pathway were consistent with previous reports of association between inflammation and elevated kynurenine in embryonic mouse brain (Goeden *et al*, 2016; Murakami *et al*, 2021). In sum, metabolomics of eCSF collected from embryos of saline-versus polyI:C-injected dams revealed differential abundance of metabolites that are part of two pathways that are relevant to the MIA model pathology – the corticosteroid and kynurenine metabolism pathways.

**Figure 3.**
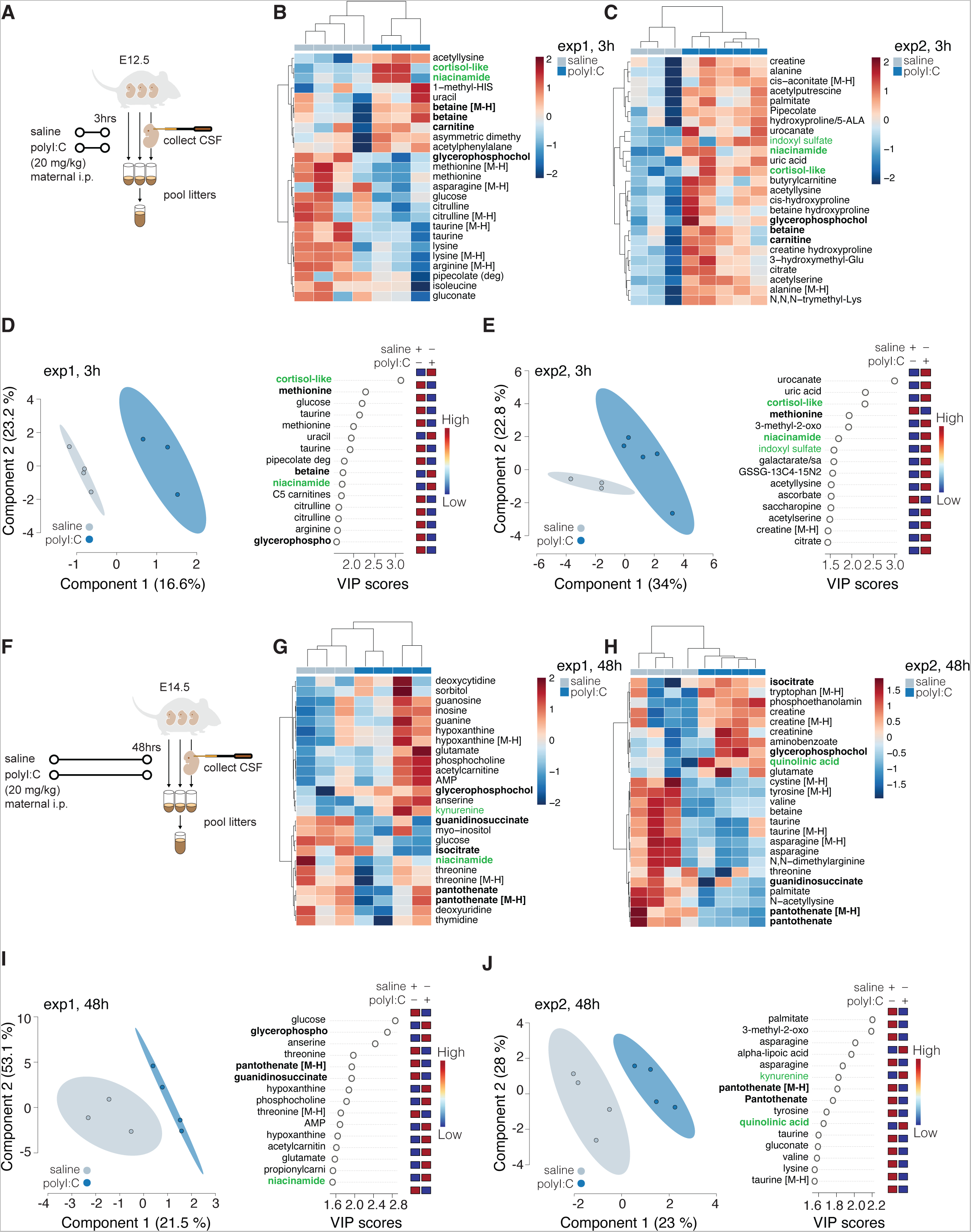
Changes observed in embryonic CSF at early (3h) or late (48h) time points after induction of inflammation in the mother by PolyI:C injection. A Schematic depicting the collection strategy of eCSF for untargeted LC-MS metabolomics 3 hours post injection with polyI:C and induction of MIA. B-E Altered metabolites 3hr post polyI:C injection. Shown are heatmaps of top 25 changed metabolites (B and C) or PLSDA and corresponding important features plots (D and E) of untargeted metabolomics analysis of polar eCSF metabolites from treatment and control samples (N>=3). Two independent experiments are presented separately (B vs C and D vs E). Metabolites of interest are highlighted in green. Metabolites overlapping between top-25 analysis and VIP analysis are in bold. Plots were generated using the online MetaboAnalyst tool, after log-transformation and Pareto scaling of the combined data from normalized positive and negative mode analysis. Level 1 (exact mass, RT match or RT and MS2 match to in-house databases) confidence metabolites were used for this analysis. F Schematic depicting the collection strategy of eCSF for untargeted LC-MS metabolomics 48 hours post injection with polyI:C and induction of MIA. G-J Same as for (B-E) except data was from samples collected 48hr post induction of MIA.

### Validation of the MIA-induced changes in corticosteroids and kynurenine pathway in eCSF

We next validated and further investigated the elevated levels of glucocorticoids and kynurenine pathway components we observed (Fig 4A, C) because these pathways are associated with inflammation and are thus relevant to the MIA model. We used the combined eCSF data from two independent experiments designated Experiments 1 and 2 (see Fig 3) at 3h following polyI:C exposure and detected two corticosterone-like molecules (Fig 4B): a cortisol-like molecule at m/z 363.2166 and a deoxycorticosterone-like molecule at m/z 344.1985, which increased approximately 2- to 5-fold in eCSF at 3h following polyI:C compared to saline control. Neither of these molecules was present in eCSF at 48h, suggesting that the eCSF metabolome can track with maternal serum metabolites. These data also suggest that the glucocorticoid aspect of the maternal inflammatory response transmitted to the embryo was largely resolved by 48h (Fig EV3D). Although we cannot currently confirm the exact identity of these corticosterone molecules, we detected elevation of multiple corticosterone metabolites (Fig 5B-D), suggesting modulation of corticosterone synthesis upon inflammation. Further investigation is required for the study of the biological role of corticosterone metabolites in neuropathology in the MIA model.

**Figure 4.**
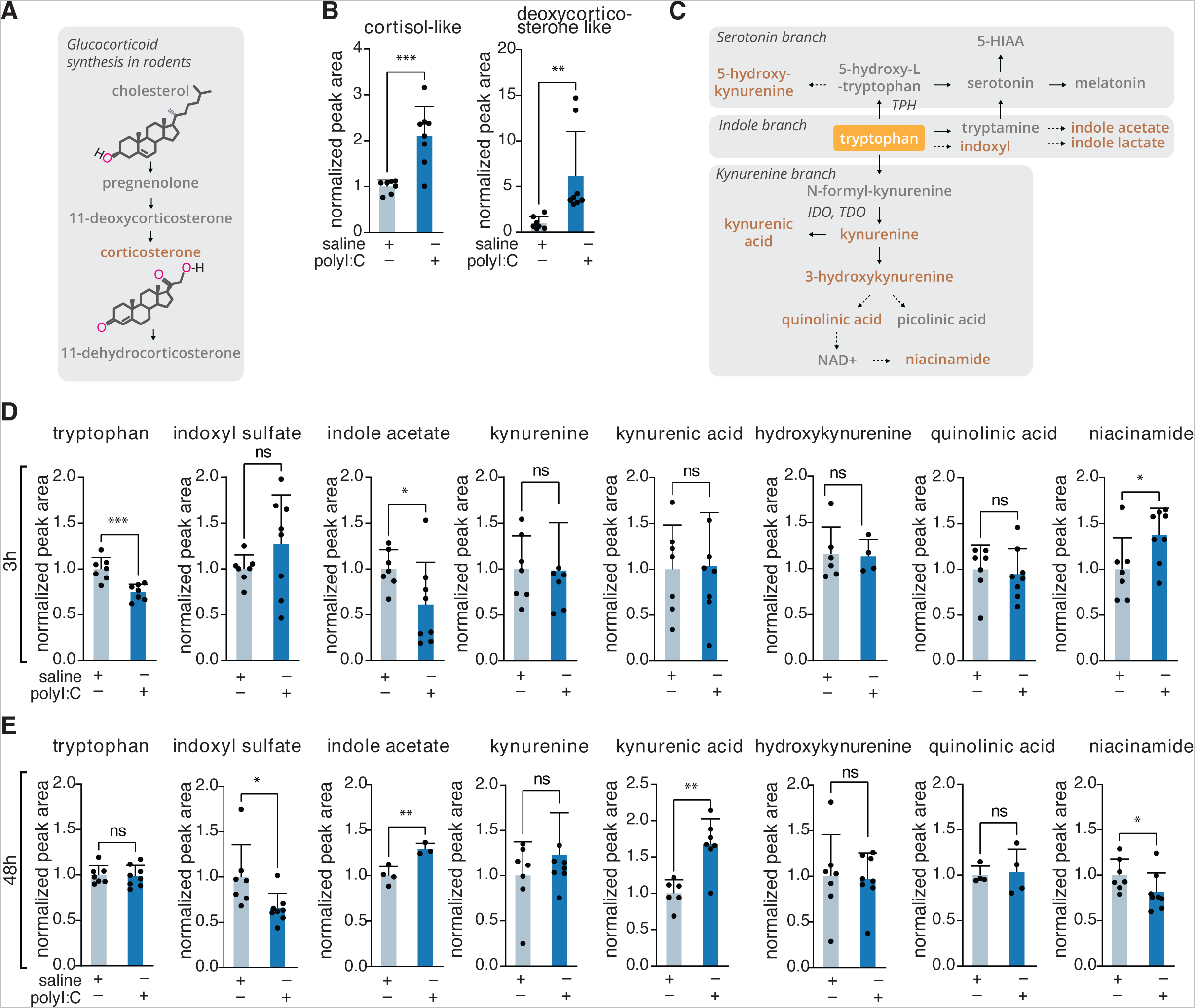
Targeted metabolomics of MIA eCSF reveals involvement of kynurenine pathway and a corticoid stress response. A Schematic of glucocorticoid synthesis in rodents. B Bar plots depicting mean +/-standard deviation of two corticosteroids identified at Level 2 confidence (including m/z match to online database with no RT) by untargeted metabolomics 3h post polyI:C injection. Two independent experiments were combined for the analysis (N>=7). C Schematic of tryptophan degradation pathways in mammals. Metabolites present in our in-house library and analyzed via targeted metabolomics approach are highlighted in dark orange. D, E Mean and standard deviation of indicated metabolites 3h (D) or 48h (E) post polyI:C injection. Data was analyzed using targeted metabolomics approach. Two independent experiments were combined for the analysis (N>=7). *= p<0.01; **= p<0.001; ***= p<0.001

**Figure 5.**
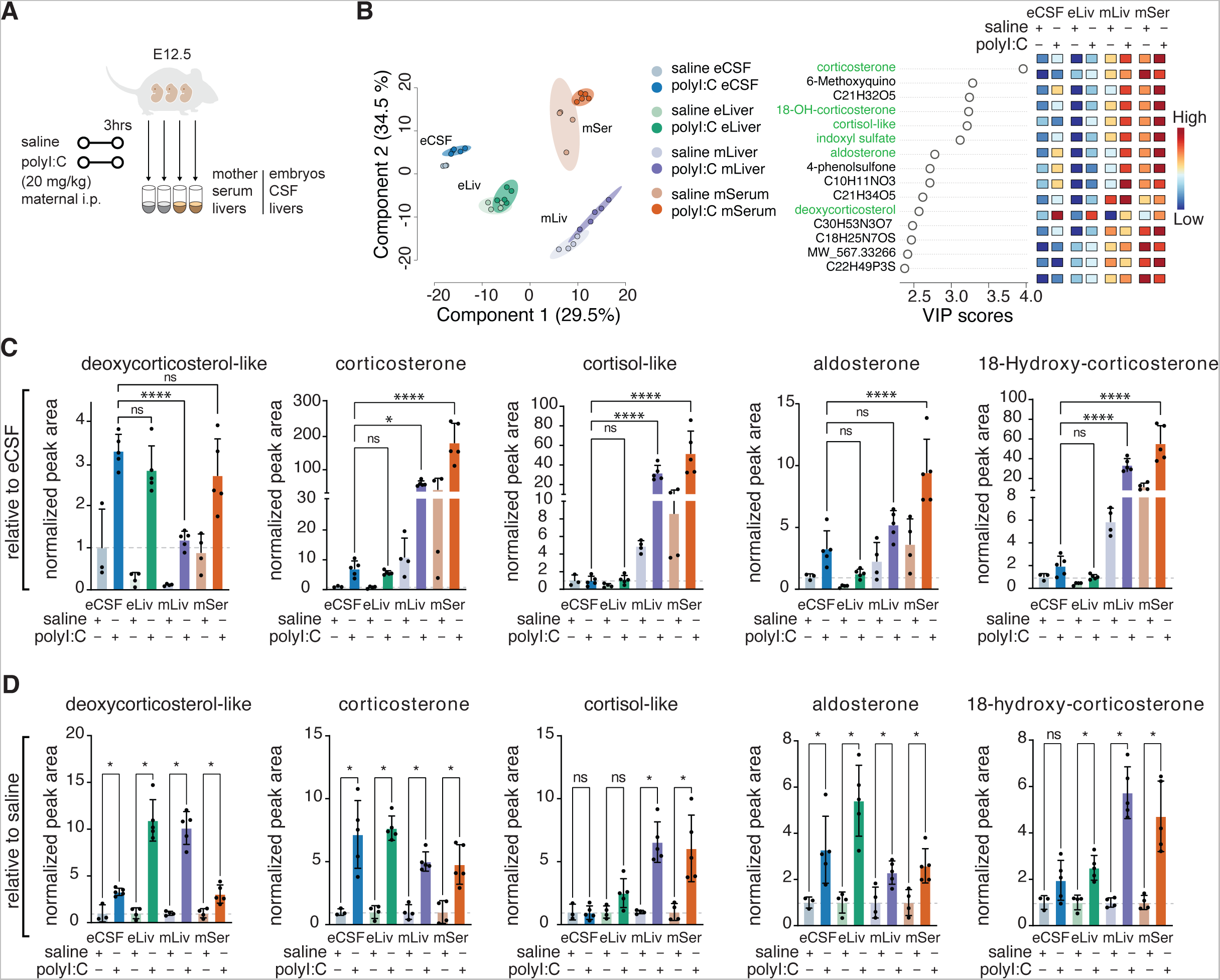
Metabolic changes as assessed by untargeted metabolomics in embryonic liver and CSF or maternal liver and serum for glucocorticoid-related intermediates. A Schematic depicting the collection strategy of embryonic CSF (eCSF) and liver (eLiv) and maternal liver (mLiv) and serum (mSer) for untargeted LC-MS metabolomics 3 hours post injection with polyI:C and induction of MIA. B Shown are PLSDA and corresponding important features plot of untargeted metabolomics analysis of polar eCSF, eLiv, mLiv and mSer metabolites from treatment and control samples at 3h post PolyI:C injection (N>=3). Plots were generated using the online MetaboAnalyst tool, after log-transformation and Pareto scaling of the combined data from normalized positive and negative-mode analysis. Level 1 to 3 (exact mass, RT match, MS^2^ match to in-house or online databases and match to eCSF CD-1-library) confidence metabolites were used for this analysis. eCSF data from experiment 2 from previous figures was from the same mice cohort and was re-analyzed for this set of graphs using CD 3.3. Metabolites of interest are highlighted in green. C, D Bar plots depicting mean and standard deviation levels of indicated metabolites post polyI:C injection for different steroids detected in our chromatography *= p<0.01; **= p<0.001; ***= p<0.001; ****= p<0.0001. Graphs are either presented relative to eCSF abundance (C) or relative to corresponding control saline sample set (D).

Next, we applied metabolite annotation verification of the kynurenine pathway (Fig 4C) using standards (Fig 4D and E). Tryptophan and indoxyl sulfate levels were reduced while niacinamide levels were increased at 3h (Fig 4D). At 48h, we observed increased kynurenic acid and indole acetate levels, while niacinamide and indoxyl sulfate levels decreased. Of note, kynurenine and quinolinic acid, which were significantly elevated in the polyI:C exposed embryos in our untargeted analysis of individual Experiments 1 and 2 (Fig 3), were not significantly higher when both experiments were combined. We optimized a separate method for detection of serotonin, picolinic acid, and 5-HIAA standards, but were unable to consistently detect these metabolites in 3-5 µl samples of eCSF, likely due to low abundance or low ionizing efficiency on the MS. These results demonstrate changes in different kynurenine pathway metabolites at the two time points analyzed and suggest alternative utilization pathways that are reported in the literature (via kynurenine or indoxyl, see Fig 4) for tryptophan degradation at 3h and 48h following polyI:C exposure.

Our data raise the question of whether the elevated kynurenine is produced locally in the embryo or originates from the pregnant dam, as kynurenine can cross the placenta (Goeden *et al*, 2017). Further, Bonnin and colleagues (2011) found that maternal tryptophan is metabolized into 5-HT in the placenta and then utilized in the fetal forebrain (Bonnin *et al*, 2011), which raises the possibility that kynurenine in the eCSF could be exogenously produced in the placenta (Goeden *et al*, 2017). The source of the kynurenine is likely to have implications on the dynamics of the accumulation and clearance of kynurenine and kynurenine-related metabolites. To address a possible maternal contribution to the elevation of kynurenine metabolites in the eCSF, and to test the dynamics of kynurenine trafficking and clearance, we next tested the effects of kynurenine supplementation to mothers (Fig EV4A). Pregnant dams received intraperitoneal (IP) injections of either saline or kynurenine (20 mg/kg). Maternal serum (mSer), embryonic serum (eSer), and eCSF were subsequently collected at different time points. Due to the inherent nature of sample collection and low volumes of fluids available at these early ages, samples were collected across different days, and replicates were obtained from independent mice. Thus, we attribute some experimental variability to technical batch differences. Nevertheless, while tryptophan levels remained unchanged (Fig EV4B), we detected clear re-normalization of kynurenine levels in maternal serum approximately 90 minutes following kynurenine supplementation (Fig EV4C). To test dynamics and distribution of kynurenine and other kynurenine pathway intermediates in the eSer and eCSF, we modeled the occurring dynamic changes assuming a linear geometric model of metabolite utilization and/or clearance (Fig EV4C-E). Kynurenine clearance or utilization seemed to follow similar dynamics to mSer, eSer, and eCSF. Thus, higher levels of kynurenine in mSer were dynamically and rapidly reflected in the levels of kynurenine in eSer and eCSF, suggesting kynurenine can quickly redistribute from the mother serum to eCSF. Similarly, kynurenic acid redistribution was not significantly different between the three biofluids but there was a trend towards a slower decay in kynurenic acid levels in eCSF, suggesting delayed clearance or utilization (Fig EV4D). We further compared the dynamic change of kynurenine, kynurenic acid and quinolinic acid within the eCSF (Fig EV4E). Here, we observed significantly different dynamics of quinolinic acid relative to kynurenine, which could be explained by a delay in the synthesis, utilization, or distribution of this metabolite in eCSF. These observations might reflect the dynamics of resolution of inflammation in eCSF and potentially explain why we see certain metabolic alterations at 48h but not at 3h post polyI:C injection in our MIA model. However, future experiments utilizing metabolic labelling will be required to expand on this finding, to distinguish between exogenous and endogenous sources of kynurenine and its intermediates, and to shed light on redistribution vs. embryonic synthesis of these metabolites.

Together, these results reveal MIA-induced changes in two metabolic pathways, corticosteroid metabolism and the kynurenine pathway. Future studies will investigate whether changes in the levels of these metabolites in the eCSF following MIA mediate, compensate, or simply reflect the severity of MIA and/or MIA-induced pathology in the embryonic brain.

### Differential dynamics of sterol and kynurenine pathway intermediates in the maternal-embryonic axis

MIA imparts systemic embryonic stress, with impact on the brain and the hematopoietic system (López *et al*, 2023; Tseng & Beaudin, 2023). At both E12.5 and E14.5, the developing embryonic hematopoietic system is located in the liver. We therefore profiled the metabolic state of the glucocorticoid and kynurenine pathways in the embryonic liver. Additionally, we analyzed the maternal liver because adult liver expresses relevant metabolic enzymes in both the glucocorticoid and the kynurenine pathways (Droin *et al*, 2021). We found differential kinetics of key metabolites between maternal and embryonic biofluids upon inflammation (Fig EV4), so we performed a side-by-side analysis between maternal serum (mSer), maternal and embryonic livers (mLiv, eLiv), and eCSF. Our data indicated that fast metabolism occurs within hours in both mothers and embryos (Fig EV4), informing relevance of the 3h time point for this analysis. We therefore applied untargeted metabolomics analysis to mSer, mLiv, eLiv and eCSF from Experiment 2 (Fig 5A) and observed a clear separation between individual sample sets (livers and biofluids) with more subtle changes between treatment and control groups (Fig 5B and Fig EV5A). Metabolites related to sterol and kynurenine pathways were observed as major drivers of the separation between all groups and among top significantly changed metabolites (Fig 5B and Fig EV5A and B). Of note, we detected metabolites from the cortisol group (Fig EV5C) that we annotate at certainty Level 3; two of these had multiple possible hits in the corticosteroid group that we labeled deoxycorticosterol-like and cortisol-like according to predicted formulas. Further work is required to delineate the exact structure and identity of these sterol intermediates and their unique roles in embryonic and maternal tissues. We cannot currently exclude that these are unique sample-specific degradation products or chromatography artifacts produced during LC-MS analysis. Nonetheless, their abundance in the analyzed tissues is consistent and we report them here alongside other glucocorticoid pathway metabolites that followed organ-specific kinetics (Fig 5C and D). A few examples of significant observations include high embryonic levels of deoxycorticosterol-like while other glucocorticoid pathway metabolites were higher in the maternal organs (Fig 5C), and all detected glucocorticoid pathway metabolites were higher in each of the analyzed organs in the polyI:C-injected compared to saline-injected dams and embryos (Fig 5D). These findings are consistent with reports that glucocorticoid receptors, glucocorticoid receptor (GR) and mineralocorticoid receptor (MR), are differentially expressed in tissues across the body and the brain (Koning *et al*, 2019). Glucocorticoid-mediated signaling could have important implications for the resolution of the propagation of inflammation in tissues across the maternal-fetal axis.

Each metabolite of the kynurenine pathway had an organ-specific distribution (Fig 6A): indoxyl sulfate was higher in maternal organs; quinolinic acid and kynurenine were higher in the embryonic organs; indole and niacinamide were higher in livers, regardless of embryonic or maternal; indole lactate was lower in the livers. Most kynurenine pathway metabolites were not significantly different in organs between saline and polyI:C injections (Fig 6B). This is aligned with our eCSF data and indicates a later response of this pathway to polyI:C (Fig 4). Tryptophan levels remained unchanged (Fig 6A, B).

**Figure 6.**
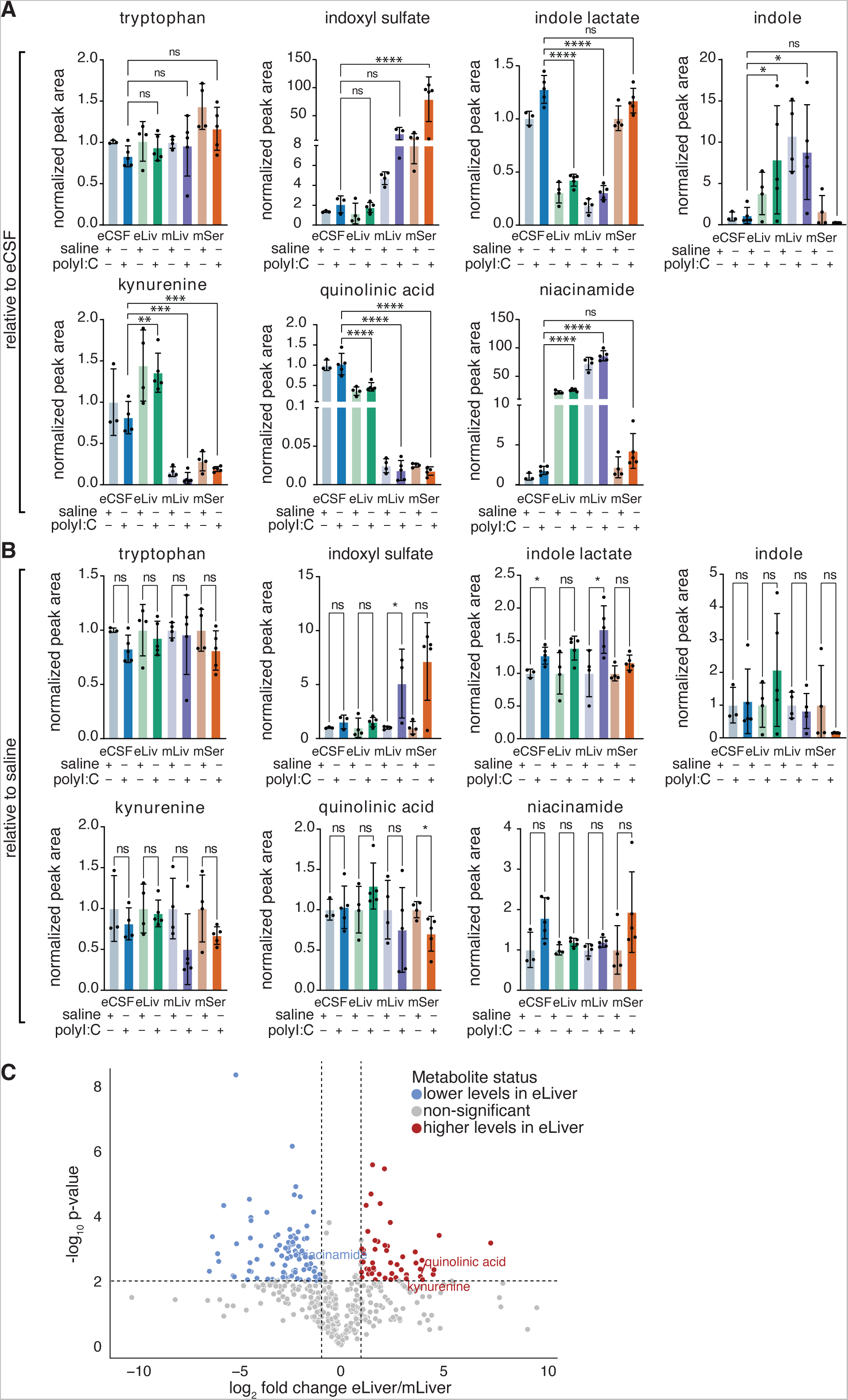
Metabolic changes as assessed by untargeted metabolomics in embryonic liver and CSF or maternal liver and serum for kynurenine pathway. A,B Bar plots depicting mean and standard deviation levels of indicated metabolites post polyI:C injection for indoxyl pathway (C and) or kynurenine pathway intermediates (E and F) by targeted metabolomics. *= p<0.01; **= p<0.001; ***= p<0.001; ****= p<0.0001. Graphs are either presented relative to eCSF abundance (A) or relative to corresponding control saline sample set (B). C Volcano plot comparing embryonic (eLiver) or maternal liver (mLiver) control samples at 3h post PolyI:C injection (N>=3). Untargeted metabolomics and Level 1 to 3 (exact mass, RT match, MS2 match to in-house or online databases and match to eCSF CD-1 library) confidence metabolites were used for this analysis. Metabolites of interest are highlighted (blue – downregulated, red – upregulated in eLiver). Only identified kynurenine pathway intermediates are annotated.

While most kynurenine pathway intermediates were not significantly altered, comparison of embryonic and maternal livers revealed that niacinamide, kynurenine and quinolinic acid were among the top significantly differentially abundant metabolites (Fig 6C), and metabolic pathways (Appendix Fig S1). Further, kynurenine and quinolinic acid were higher in eLiv samples relative to mLiv levels, while niacinamide was higher in mLiv compared to eLiv (Fig 6A and C). Collectively, our results indicate distinct dysregulation of the kynurenine pathway in the mother and embryo in the MIA model. This effect could result from maternal inflammation or an embryonic response to maternal stress, since embryonic changes can reflect embryonic metabolism or an overflow of maternal metabolites, and both possibilities likely respond to maternal inflammation.

## Discussion

CSF composition, including proteins and small molecules, is critical for supporting brain development (Lehtinen *et al*, 2011), and changes in CSF composition during development may contribute to neuropathologies (Zappaterra & Lehtinen, 2012). (Lehtinen *et al*, 2011)(Zappaterra & Lehtinen, 2012)Profiling eCSF composition can reveal abnormalities. For example, we previously identified pro-inflammatory peptides (e.g., cytokines) in eCSF following MIA (Cui *et al*, 2020). However, the developing eCSF is currently underutilized for uncovering potential effectors of CNS development and disease. There are available mouse models of pathologic conditions that often lead to neurodevelopmental disorders in human, such as the MIA model. However, in part due to challenges in obtaining sufficient quantities of mouse eCSF, broad, comprehensive profiling of mouse eCSF has been extremely challenging. To date, only one metabolite profiling study of the eCSF has been published (Requena-Jimenez *et al*, 2021). No metabolite profiling of MIA CSF has been previously published, even though the metabolite content of the CSF is likely critical for MIA pathophysiology.

To gain a more complete picture of eCSF metabolite content, we developed a pipeline to obtain metabolite profiling data from eCSF and generated a small molecule database of healthy CD-1 eCSF on embryonic day E14.5. This database can be used as a refence library of the eCSF metabolite content and is therefore a new resource for researchers in the fields of neurobiology and development. Further, we utilized the eCSF metabolite library dataset as a reference library in a comparison of the eCSF metabolome between two mouse strains (C57BL/6 and CD-1), two developmental days (E12.5 and E14.5), and in normal vs. pathologic condition (MIA induced by polyI:C exposure).

We found evidence that two developmentally essential metabolic pathways were altered by MIA in eCSF: the glucocorticoid and kynurenine pathways. First, corticosterone was elevated 3hrs following MIA, a time prior to the appearance of pro-inflammatory cytokines in the eCSF but coinciding with cytokines and corticosterone elevations in the maternal plasma (Smith *et al*, 2007; Choi *et al*, 2016; Meyer *et al*, 2006; Estevez *et al*, 2020). Corticosterone can cross the placenta; therefore, maternal serum is a likely source of eCSF-corticosterone. (Smith *et al*, 2007; Choi *et al*, 2016; Meyer *et al*, 2006; Estevez *et al*, 2020)Once in the brain, corticosterone is a potential candidate for mediating the effects of MIA on embryonic brain development by multiple mechanisms, such as accelerating neural progenitor proliferation (Tsiarli *et al*, 2017, 2013). Glucocorticoids can signal through glucocorticoid receptor (GR) and mineralocorticoid receptor (MR), both of which are differentially expressed in tissues across the body and the brain (Koning *et al*, 2019; Russell & Lightman, 2019) and are reported to have opposing roles during inflammation and stress (Koning *et al*, 2019; Chantong *et al*, 2012). Thus, future studies will be necessary to determine if CSF-corticosterone directly impacts forebrain neurogenesis in the context of MIA.

A second set of metabolites that we found to be affected in eCSF following MIA originate from tryptophan catabolism and includes kynurenine, kynurenic acid and quinolinic acid (Goeden *et al*, 2016). The kynurenine pathway was altered 48 hours following MIA, a time that coincides with elevated cytokine levels in CSF (Cui *et al*, 2020) and brain (Meyer *et al*, 2006; Choi *et al*, 2016), which are indicative of embryonic inflammation. Whether the source of eCSF kynurenine is maternal or embryonic is currently unclear. Kynurenine can pass from maternal plasma (Goeden *et al*, 2017) and be metabolized by the placenta (Goeden *et al*, 2016), and even more proximally - by the embryonic liver (Mu *et al*, 2020). eCSF kynurenine can also be metabolized locally in the embryonic CNS (Savitz, 2020). (Goeden *et al*, 2017)(Goeden *et al*, 2016)(Mu *et al*, 2020)(Savitz, 2020)Once in the eCSF, changes in metabolites of the kynurenine pathway have the potential to mediate MIA effects on development. For example, kynurenic acid and quinolinic acid are both ligands for the N-methyl d-aspartate receptor, which modulates the proliferation and maturation of glutamatergic cells (Toriumi *et al*, 2012). These cells are affected in MIA-induced cerebral cortical disorganization models (Hsiao & Patterson, 2011; Choi *et al*, 2016; Cui *et al*, 2020), suggesting a potential link between metabolites of the kynurenine pathway and disrupted cerebral cortical development.

Our findings raise intriguing questions regarding the routes by which peripheral metabolites gain access to the embryonic CSF and brain. Most likely, the main routes for peripheral metabolite transfer are through the choroid plexus (Fame & Lehtinen, 2020) and the blood-brain barrier (BBB); neither are fully developed in the embryo (O’Brown *et al*, 2018). Of note, the embryonic barrier mechanisms are differentially regulated compared to adult BBB (Møllgård *et al*, 2017). Additionally, the developing circumventricular organs could also provide a plausible route for peripheral metabolite molecules to enter the CSF (Kiecker, 2018), as this has been described to occur in adult brain (Skovbjerg *et al*, 2023; Broadwell & Brightman, 1976).

Our findings begin to shed light on important questions regarding (1) the kinetics of a developing embryo sensing and acquiring maternal illness and (2) how long a maternal infection persists in the developing embryo, despite maternal recovery. Indeed, our data suggest that temporal regulation of maternal-fetal illness can be evaluated by different metabolite pathways. The corticosterone pathway provides an early indication of maternal illness in the developing embryo whereas other metrics of inflammation, including kynurenine, and previously published cytokine profiles (Cui *et al*, 2020), contribute to later inflammatory signatures of CSF. Notably, many metabolites detected at E12.5 were significantly less abundant at E14.5 (Fig 2D). Future analyses will determine if the apparently reduced abundance was due to further metabolism of pathway intermediates, a dilution effect due to increased embryo growth / CSF volume, or some level of CSF replacement / turnover already present at this early age. Collectively, our findings raise intriguing questions about target cells in the developing brain and long-term consequences on neural circuitry and lifelong functions.

Importantly, the polyI:C MIA model represents just one available model for investigating the consequences of maternal illness on the developing brain. The metabolic maternal-fetal axis is sensitive to perturbation from many types of challenges including other forms of maternal illness, diet, substance abuse, and environmental teratogens, all of which are associated with manifestations of later onset neurologic and psychiatric conditions. Thus, our resource takes a first step towards elucidating how the eCSF is altered in MIA and provides a path forward for investigating whether similar mechanisms and kinetics generalize beyond MIA to other conditions as well.

### Limitations and future directions

A limitation of our study stem from the use of polyI:C to model MIA (Kowash *et al*, 2019; Meyer *et al*, 2006). To the best of our abilities, we have controlled for potential biological noise due to inter- and intra-bottle variability by tracking reagent lot number, verifying molecular weight, and tracking endotoxin levels, as recommended in Kowash et al., 2019 (Kowash *et al*, 2019). Yet, biological noise between and within our experiments was observed and this noise confined the statistical power of our study. In future studies it will be important to determine the degree to which polyI:C-associated metabolite changes in the eCSF compare with other models of MIA including other forms of maternal illness or environmental teratogens and pollutants (Bucknor *et al*, 2022; Block *et al*, 2022).

At the metabolomics analysis level, our study was constrained by the low volume of eCSF, dictating limitations of sample re-runs on the LC-MS for quality assurance or analysis with multiple chromatography methods. Therefore, our study focused on the profiling of polar metabolites. Future studies can improve the eCSF metabolomics dataset through expansion of chromatographic conditions, to allow characterization of eCSF to compound groups that include non-polar compounds, fatty acids, sterols, lipids, hormones, and more.

Our database is annotated at different levels of certainty scored in accordance with the metabolomics community recommendations (Salek *et al*, 2013; Fiehn *et al*, 2007; Sumner *et al*, 2007). A fraction of entries (LC-MS features) in our database are only specified by a chemical formula or a measured exact mass (m/z) and do not match internal or external databases. Thus, we did not score these entries highly in our identification certainty scoring system. These unidentified metabolites are potentially a valuable recourse to the community because: (1) we have performed extensive filtering steps to ensure that they originate from eCSF rather than represent contaminants or background, (2) they can be identified chemically, as they are present in appropriate quantities in eCSF to be well detected by LC-MS, (3) they can serve as biomarkers of neuropathology that stems from MIA, and (4) they can generate testable hypotheses and help with further characterization of the eCSF’s biochemical properties, normal function, or disease states. However, as much as many of these unidentified metabolites are potentially novel CSF-specific compounds and should not be disregarded as biologically irrelevant, these lowly-scored metabolites might also be the result of the formation of adducts or fragments during analysis, and therefore may be redundant, secondary products of known compounds. Further extensive *in silico* and *in vitro* characterization will be needed to accurately characterize true novel compounds.

In the future, our embryonic mouse CSF metabolomic resource would be instrumental in research exploring how different brain regions respond to MIA at various stages of development. Our data complement studies like the recent one describing MIA’s impact on the developing hippocampus, which found key differentially abundant metabolites (e.g., amino acid metabolism, valine, and leucine biosynthesis, etc.)(Southey *et al*, 2023). Together with these existing data, our resource will contribute to a more mechanistic understanding of MIA relevant processes. In a separate approach, our resource can be used to generate hypotheses for how glutamate, a neurotransmitter already implicated in MIA neuropathology (Southey *et al*, 2023; Murakami *et al*, 2023) might exerts its effects. Our dataset could also help define metabolites with temporally restricted profiles (e.g., E12.5 only or E14.5 only), similarly to certain ions (Xu *et al*, 2021), thus guiding the formulation of an age-specific embryonic CSF for future *ex vivo* studies.

### Summary

Childhood brain development relies on appropriate *in utero* maternal-fetal health when the blueprint for the maturing brain is established (Wilde *et al*, 2014; Lu-Culligan & Iwasaki, 2020). Our data provide a unique perspective on this critical blueprint in the form of extensive metabolite profiling of select fluids and tissues across the maternal-fetal axis, with a focus on the eCSF. Comparative untargeted and targeted metabolomics, as reported here, have the potential to illuminate an array of future research directions ranging from identification of biomarkers and improved diagnostics to possible targeted therapeutics that will ameliorate altered neurodevelopment.

## Author contributions

Conceptualization: B.P., T.E.L., N.K., and M.K.L.; Formal analysis: B.P., T.E.L., A.C., and A.M.; Investigation: B.P., T.E.L., and A.C.; Methodology: B.P., T.E.L., and A.C.; Writing original draft: B.P., T.E.L., N.K., and M.K.L.; N.K. and M.K.L. provided research support; Writing review & editing: all authors. All authors have read and agreed to the published version of the manuscript.

## Institutional Review Board Statement

This research complies with all relevant ethical regulations and was approved by Boston Children’s Hospital IACUC and Boston Children’s Hospital Institutional Biosafety Committee. The Boston Children’s Hospital IACUC approved all experiments involving mice in this study.

## Data Availability Statement

Data associated with this study will be made available at MetabolomicsWorkbench: https://www.metabolomicsworkbench.org/

## Supporting information

supplementary_materials_MIA

## Acknowledgments

We thank all members of the Kanarek and Lehtinen labs for their advice and help. We thank Nancy Chamberlin, Peng Wang, Amy Yu, Huixin Xu, Ryann Fame, Chris Beecher, and Felice de Jong for critical reading of the manuscript. We thank Chris Beecher and Felice de Jong for valuable discussions and input on ClusterFiner software and Eric Tague for help on CompoundDiscoverer.

## Methods and Protocols

### Animals

All mouse work was performed in accordance with the Institutional Animal Care and Use Committees (IACUC) and relevant guidelines of Boston Children’s Hospital. Embryonic day (E) 14.5 embryos from time-pregnant CD-1 (ICR) dams were purchased from Charles River Laboratories (CRL) for the baseline conditions metabolomics database and maternal kynurenine injection time-course experiments. E12.5 embryos from timed-pregnant C57BL/6 dams were purchased from CRL for maternal immune activation metabolomics experiments. For timed pregnancies, females were checked for the presence of plugs, and the date of the plug was noted as embryonic day 0.5 (E0.5). All animals were housed under 12hr/12hr day night cycle with access to standard chow and water *ad libitum*.

### Embryonic and maternal body fluid and tissue collection and analysis

Embryonic CSF (eCSF) was collected by inserting a glass capillary into the cisterna magna and processed as described (Zappaterra *et al*, 2013). Each CSF sample was pooled across litters, purity was verified visually with a dissection microscope, and then frozen on dry ice. Embryonic blood was collected by glass capillary following a neck artery nick and pooled across litters. Maternal blood was collected either by tail vein nick for non-terminal timepoints or intracardiac puncture for terminal timepoints. Serum samples were analyzed following coagulation, centrifugation, and then immediately put on dry ice. Embryonic and maternal livers were dissected and immediately transferred to dry ice.

### Maternal immune activation

On E12.5, pregnant dams received a single dose (20 mg/kg, i.p.) of polyinosinic-polycytidylic acid (polyI:C, Sigma Aldrich) or sterile saline as vehicle control. To confirm maternal immune activation, maternal blood was collected at 3hrs following maternal polyI:C injection by either tail-nick or intracardiac puncture. After maternal blood samples coagulated and centrifuged, serum samples were analyzed by LEGENDplex Mouse Inflammation Panel (13-plex with V-bottom Plate) FACS-ELISA (BioLegend).

### Sample preparation for LC-MS analysis of polar metabolites from CSF, plasma, and liver from embryonic and adult mice

We used fresh frozen tissues or processed (see method above) CSF and plasma. Per condition, 5-10 µl embryonic CSF, 5 µl plasma or a grain-of-rice sized sample of maternal and embryonic livers were extracted. Tissues were homogenized with a handheld homogenizer (Sigma, Z359971) and briefly sonicated.. 300µl of extraction solvent (80% LC/MS-grade methanol, 20% 25 mM Ammonium Acetate and 2.5 mM Na-Ascorbate prepared in LC/MS water and supplemented with isotopically labeled internal standards (17 amino acids and isotopically labeled reduced glutathione, Cambridge Isotope Laboratories, MSK-A2-1.2 and CNLM-6245-10)) was used per sample. After centrifugation for 10 min at maximum speed on a benchtop centrifuge (Eppendorf) the cleared supernatant was dried using a nitrogen dryer (ThermoFisher Scientific, TS-18826) and reconstituted in 17 µl (eCSF) or 25ul (plasma, tissues) “QReSS” water (supplemented with QReSS, Cambridge Isotope Laboratories, MSK-QRESS-KIT) by brief vortexing. Extracted metabolites were spun again and the cleared supernatant was transferred to LC-MS micro vials. A small amount of each sample was pooled and serially diluted 3 and 10-fold for quality controls (pool-QC) throughout the run of each batch. About ½ of eCSF (7µl), 1/25^th^ of plasma (1µl), and 1/25^th^ of tissue (1µl) were injected onto a chromatographic column and subsequently analyzed using LCMS.

### Chromatographic conditions for LC-MS

ZIC-pHILIC chromatography was used for polar metabolites: an appropriate volume of reconstituted sample was injected into a ZIC-pHILIC 150 × 2.1 mm (5 µm particle size) column (EMD Millipore) operated on a Vanquish™ Flex UHPLC Systems (Thermo Fisher Scientific, San Jose, CA). Chromatographic separation was achieved under the following conditions: buffer A was acetonitrile; buffer B was 20 mM ammonium carbonate, 0.1% ammonium hydroxide. Gradient conditions were: linear gradient from 20% to 80% B; 20–20.5 min: from 80% to 20% B; 20.5–28 min: hold at 20% B. The column oven and autosampler tray were held at 25 °C and 4 °C, respectively. Chromatographic performance was quality controlled using a mixture of unlabeled standard amino acids and a mixture of chemically diverse compounds (Cambridge Isotope Laboratories, MSK-A2-US-1.2 and MSK-QRESS-US-KIT) with 1µl of each injected before or after every run on our HILIC method.

### MS data acquisition conditions for targeted or untargeted analysis of polar metabolites

MS data acquisition was performed using a QExactive benchtop orbitrap mass spectrometer equipped with an Ion Max source and a HESI II probe (Thermo Fisher Scientific, San Jose, CA) and was performed in positive and negative ionization mode in a range of m/z= 70–1000, with the resolution set at 70,000, the AGC target at 1×10^6^, and the maximum injection time (Max IT) at 20 msec. Optimal HESI conditions were determined utilizing a mix of chemical standards: Sheath gas flow rate: 35; Aug gas flow rate: 8; Sweet gas flow rate: 1; Spray voltage: 3.5kV (pos), 2.8kV (neg); Capillary temperature: 320°C; S-lens RF: 50; Aux gas heater temperature: 350 °C. For untargeted analysis, additional Top8 method (see below) was employed in either positive or negative mode with ddMS settings as: the resolution set at 17,500, the AGC target at 1×10^5^, the maximum IT at 50 msec, isolation window at 1.0 m/z, and stepped NCE at 20,40 and 80. As our software/hardware did not allow for the widely employed AquireX assisted MS2 inclusion/exclusion list data collection, we performed repeated data collections with the following logic: At the end of each experimental batch spectral data was collected in both positive and negative mode on additional injections of the pool-QC. We collected spectral data in a data-dependent acquisition mode (DDA) such that for every MS^1^-level scan the top 8 most abundant ions were picked for fragmentation (Top8-level experiment), an exclusion window was set to 6 seconds, such that ions picked for fragmentation would be put on an exclusion list for 6 seconds before they could be picked for fragmentation again. This allowed for the collection of about 300-500 individual ion spectral scans per sample injection. Then, MS^1^ and MS^2^ level data were quickly analyzed using CompoundDiscoverer and detected features were used to create an inclusion and exclusion list (per polarity) that ensured maximal inclusion of interesting features (features with signals above mock level and with no MS^2^ spectra collected) and maximal exclusion of non-interesting features (features present in mock samples for which MS^2^ spectra was already collected). This strategy was then performed iteratively up to 4 times. We noted an incremental increase in the number of features with associated spectra generated with this strategy, however we note that this strategy did not perform as well as repoted from an AquireX type of experiment.

### Generation of targeted metabolomics libraries (in-house MS^1^ and MS^2^-level databases)

Two separate MS^1^ and MS^2^-level libraries were generated. A set of standards premixed in 7 individual pools were run on our LC-MS HILIC method. Information on this pool set has been previously published (Petrova *et al*, 2021; Sullivan *et al*, 2019). Briefly, we first collected MS^1^ level information (retention time and peak shape and quality were noted, see Table EV1). To generate the MS^2^-level database, each pool set was injected twice (for positive and negative mode), and specific inclusion lists were preset for a PRM-type of experiment. This inclusion list was based on the already determined retention times and was focused on compounds with peaks of minimum acceptable quality. Thus, high quality MS^2^ data were collected per individual metabolite and MS^2^ spectra were averaged out from three different HCD cell energies: 20, 40, 80 NCE. We named this database “HILIC_all” (Dataset EV1). The second library set was based on the MSMLS library of compounds (MSMLS, IROA Technologies’), which we named “MSMLS_HILIC”. Compounds were solubilized and pooled based on manufacturer’s recommendations. Each pool was run as an individual injection and MS^1^ information (retention time and peak shape and quality) was noted. (Table EV2). Then, spectral MS^2^ information was collected at both ionization modes as above with the exception that individual spectra were not averaged but collected at the three different HDC energies independently (Dataset EV2). All spectra were manually inspected and compared to known and well characterized spectral databases (MONA and mzCloud^TM^). Most of the spectra included in our library were thus verified. Where no match was found but the spectrum was of high quality, we included it in the library and added a note in our metadata file (Dataset EV3 (HILIC_all) and EV4 (MSMLS_HILIC)). Few metabolites were also run from individual wells, to try and obtain better signal-to-noise ratios. Overall, our database included: HILIC-all MS^1^ – 261 compounds, MS^2^ – 144 compounds; MSMLS_HILIC MS^1^ – 236 compounds; MS^2^ – 307 compounds. The overlap between the two MS^1^ databases was about 100 compounds.

### Targeted metabolomics data analysis

Relative quantification of polar metabolites was performed with TraceFinder 5.1 (Thermo Fisher Scientific, Waltham, MA, USA) using a 5 ppm mass tolerance and referencing an in-house library of chemical standards (HILIC_all and MSMLS_HLIC). We routinely queried about 257 compounds (37 internal standards and 220 metabolites). Pooled samples and fractional dilutions were prepared as quality controls and injected at the beginning and end of each run. In addition, pooled samples were interspersed throughout the run to control for technical drift in signal quality as well as to serve to assess the coefficient of variability (CV) for each metabolite. Data from TraceFinder were further consolidated and normalized with an in-house R script: (https://github.com/FrozenGas/KanarekLabTraceFinderRScripts/blob/main/MS_data_script_v2.4_20221018.R). Briefly, this script performs normalization and quality control steps: 1) extracts and combines the peak areas from TraceFinder output .csvs; 2) calculates and normalizes to an averaged factor from all mean-centered chromatographic peak areas of isotopically labeled amino acid and QReSS internal standards within each sample; 3) filters out low-quality metabolites based on user entered cut-offs calculated from pool reinjections and pool dilutions; 4) calculates and normalizes for biological material amounts based on the total integrated peak area values of high-confidence metabolites. In this study, the linear correlation between the dilution factor and the peak area cut-offs are set to RSQ>0.95 and the coefficient of variation (CV) < 30%. Finally, data were log transformed and Pareto scaled within the MetaboAnalyst-based statistical analysis platform (Xia *et al*, 2015) to generate PCA, PLSDA, volcano plots, and heatmaps. Individual metabolite bar plots and statistics were calculated in Excel and GraphPad Prism.

### Untargeted metabolomics data analysis using CompoundDiscoverer (mock-filtering mode)

Relative quantification for untargeted polar metabolomics was performed with Compound discoverer 3.3. A general workflow was built to best suit our polar metabolomics LC-MS method (details on each parameter are given in Dataset EV5). Positive and negative modes were analyzed separately. Both MS^1^ and MS^2^-level in-house databases were used (HILIC_all and MSMLS_HLIC). For the analysis of MIA samples, additionally MS^1^ and MS^2^-level CD-1-DBs were used (in positive and negative modes, see above). Filtering steps were performed based on peak noise levels, ppm error, formula annotation, and the relative abundance of the integrated peak area in true sample compered to mock samples, where features with +>3-fold higher in samples were retained. Normalization steps were performed based on a “scaling factor”: a per sample normalizer. This scaling factor was determined per experiment or sample comparison: each sample was analyzed by targeted metabolomics first, and the scaling factor corresponded to the biological material normalizer from total integrated peak area of mean-centered values from high-confidence metabolites (see above). This normalizer rarely differed more than 2-fold within a sample type. When experiments from different experimental days were compared (such as CD-1 to C57BL/6 comparison, as in Figure 2), untargeted analysis was performed on the combined samples and data were independently curated. This ensured adequate chromatographic retention time correction and alignment and thus proper feature alignment, identification, and annotation. Within our HILIC chromatography, we rarely observe chromatographic shift larger that 1 minute and retention times drifts larger than 40 seconds. We further observed that larger threshold for retention time shift led to mis-annotation for our in-house library compound databases (for example betaine vs valine or alanine vs beta-alanine). Therefore, these parameters were set as limits for retention time correction and matching to internal databases. Post CompoundDiscoverer analysis included merging of negative and positive mode data and statistical analysis using the MetaboAnalyst platform. As compound annotation at levels higher than Level 1 include a significant level of uncertainty, we decided not to merge features based on annotation name but rather to retain information from both positive and negative mode analyzed data (thus multiple compounds are annotation as either “Metabolite_name_[M-H]” or “Metabolite_name” for negative or positive mode respectively, for example Figure 2E). Finally, data were log transformed and Pareto scaled within the MetaboAnalyst-based statistical analysis platform (Xia *et al*, 2015) to generate PCA, PLSDA, volcano plots, and heatmaps. Individual metabolite bar plots and statistics were calculated in Excel and GraphPad Prism. Pathway enrichment was similarly performed within MetaboAnalyst following the above data transformation steps.

### Untargeted metabolomics data analysis using ClusterFinder and IROA credentialing (IROAmode)

All steps for sample preparation for IROA-assisted metabolomics were as described above except for the final resuspension step, which was performed using the IROA provided Internal Standard (IS) (95% ^13^C-labeled, IROA® TruQuant Yeast Extract Semi-targeted QC Workflow, IROA Technologies™). Relative quantification based on IROA for untargeted polar metabolomics was performed with ClusterFinder 4.2.23. We initially determined that 7ul of eCSF reconstituted in 20µl IS led to optimal signal intensity correspondence between labeled and unlabeled isotopologues (see Figure S1, step 1). Default parameters were set for each search (targeted or untargeted) except for the minimal signal intensity threshold which was set to 10.000. Additionally, we integrated our targeted metabolomics MS^1^-level databases into ClusterFinder and thus performed a secondary level of annotation within the software. Retention time correction and peak integration was performed within the software and was not further curated.

### Generation of untargeted metabolomics CD-1 library (CD-1-DB MS^1^ and MS2-level database)

Embryonic CSF from E14.5 embryos was collected as described above. 7 mice were used and CSF from all embryos was pooled per mouse. For each replicate, 7µl were used for IROA-assisted analysis and 7µl for mock-filtering analysis. As IROA analysis did not yield satisfactory annotation we only generated eCSF-specific metabolite databases from mock-filtering strategy. Within CompoundDiscoverer, we followed the steps outlined above. We extensively curated each feature. Features were given identifying tags that would help organize certainty of identification and annotation (Level 1-4). All features with matches to our in-house database were confirmed after manual inspection and any double annotations removed (as well as renamed and specifically tagged) if retention times were not ideal match. Both MS^1^ and MS^2^ level databases were generated within CompoundDiscoverer on curated data. After export of MS^2^ library to mzVault, the CD-1-eCSF database was further manually curated, where low quality spectra were removed. Of note – most features are represented by a single collected spectrum.

