## supplementary_materials_MIA for "Metabolomics of Mouse Embryonic CSF Following Maternal Immune Activation"

**Author Affiliations**
1. Department of Pathology, Boston Children's Hospital, Boston, MA 02115, USA;
2. Harvard Medical School, Boston, MA, USA 02115;
3. Graduate Program in Biological and Biomedical Sciences, Harvard Medical School, Boston, MA 02115, USA; Boston, MA 02115;
4. Broad Institute of Harvard and Massachusetts Institute of Technology, Cambridge, MA 02142, USA.
5. IROA Technologies[TM] LLC, Chapel Hill, NC, USA
6 Harvard-MIT Division of Health Sciences and Technology, Massachusetts Institute of Technology, Cambridge, MA 02139

Figure EV1. Schematic representation of the steps involved in IROA-assisted untargeted LC-MS analysis as performed within the ClusterFinder analysis software. Left panel: As a first step, different amounts of mouse CSF (depicted in grey) were mixed with fixed amount of IROA internal standards (IS, at 5:95 unlabeled ($^{12}C$) and $^{13}C$-labeled metabolome, as depicted in blue). This determined the optimal ratio (4 µl eCSF to 20 µl IS) – where majority of detected features were observed at one-to-one ratio. Right panel: Step two (experiment presented in Fig 1D) generated a database of compounds based on untargeted LC-MS metabolomics using IROA LTRS standard: a 1/1 mix of 95:5/5:95 unlabeled ($^{12}C$) and $^{13}C$-labeled reference metabolome ($^{12}C/^{13}C$ reference). Only features with credentialed signal (that is comprised of labeled and unlabeled isotopologues) were considered. Then samples prepared using the optimal ratio of IS to CSF were queried in a quasi-targeted manner – referencing the LTRS database. Features were further annotated using in-house databases. Phenylalanine is given as an example of credentialed feature, metabolite x – as an example of non-credentialed (orphan m/z) signal.

Figure EV2

A, B Shown are heatmap (A)and PCA (B) plots of CD-1 and C57BL/6 eCSF untargeted metabolomics samples at the corresponding embryonic stage (E14.5). (N>=3). Plots were generated using the online MetaboAnalyst tool, after log transformation and Pareto scaling of the combined data from normalized positive and negative-mode analysis. Compounds present in the in-house and CD-1-based embryonic CSF libraries (at $MS^1$-level) were used for the heatmap comparison (A) while level 1-3 confidence metabolites (including matches to the CD-1 database presented in Figure 1) were used for the PCA in B. In B, also mock samples were included for comparison.

Figure EV3. **Application of eCSF library to maternal immune activation (MIA) model**.

A,B Schematic (A) and ELISA (B) assessing the inflammatory response in the maternal serum cytokines 3h post injection of maternal polyI:C. This experiment corresponds to experiment 1 at 48h post polyI:C (Figure 3 G and I). Graphs represent mean +/- standard deviation. Statistical testing was based on unpaired t test for each cytokine, where *= $p<0.01$; **= $p<0.001$; ***= $p<0.001$; n.s.=not significant.

C,D Schematic (C) and ELISA (D) assessing the inflammatory response in the maternal serum cytokines 48h post injection of maternal polyI:C. This experiment corresponds to experiment 2 at 48h post polyI:C (Figure 3H and J). Graphs represent mean +/- standard deviation. Statistical testing was based on unpaired t test for each cytokine, where *= $p<0.01$; **= $p<0.001$; ***= $p<0.001$; n.s.= not significant.

E,F Volcano plots of untargeted metabolomics analysis of polar eCSF metabolites from treatment and controls at two time points: 3h (E, experiment 1 and 2 presented separately) or 48h (F, experiment 1 and 2 presented separately) post polyI:C injection (N>=3). Significantly changed metabolites are highlighted, only metabolites present in our in-house database are annotated. Red – upregulated, blue- down regulated, green – metabolites of interest. Data was analyzed using the online MetaboAnalyst tool, after log transformation and Pareto scaling of normalized positive and negative-mode combined datasets. Level 1-3 (including matches to the CD-1 database presented in Figure 1) confidence metabolites were used for this analysis.

Figure EV4. **Perturbation of kynurenine levels in the mother translate to measurable changes of kynurenine pathway intermediates in embryo**.

A Schematic depicting the IP-injection strategy for kynurenine-levels perturbation. Several time points were collected, time point zero was from mothers injected with saline. Embryonic CSF (eCSF) and serum (eSer) and maternal serum (mSer) were collected for targeted LC-MS metabolomics. CSF from embryos from a single mother was pooled for analysis.

B Mean and standard deviation of tryptophan levels in indicated biofluids at different time points upon injection of kynurenine in the mother. Each replicate was an independent pregnant dam or a pooled litter of eSer or eCSF and data from three sample collections (except time point 15min, which was only performed twice) was normalized to saline control dams.

C-D Left panels: as in B but for kynurenine(C), kynurenic acid (D). Right panels: corresponding by metabolite trajectory plots for relative levels in indicated biofluids. Statistics model testing was based on logarithmic decay where slopes of log-transformed data were compared. Individual p-values are depicted.

E As in C and D but relative levels of kynurenine, kynurenic acid, and quinolinic acid in eCSF are compared across different time points post injection of kynurenine in the mother.

Figure EV5. **Assessment of metabolic changes in embryonic liver and CSF or maternal liver and serum by untargeted metabolomics.**

A Schematic of glucocorticoid synthesis in rodents. Metabolites annotated in our untargeted analysis are highlighted in dark orange.

B Heatmap of Top50 changed metabolites in embryonic liver (eLiv) and eCSF or maternal liver (mLiv) and serum (mSer) by untargeted metabolomics. Level 1-3 (including matches to the CD-1 database presented in Figure 1) confidence metabolites were used for this analysis. Data was analyzed using the online MetaboAnalyst tool, after log transformation and Pareto scaling of normalized positive and negative-mode combined datasets.

B Volcano plot, depicting a comparison between polar metabolites detected by untargeted metabolomics from eLiv and eCSF or mLiv and mSer upon polyI:C injection. Level 1-3 confidence metabolites were used for this analysis. Significantly changed metabolites are highlighted, only metabolites present in our in-house database are annotated on the graph. Red – upregulated, blue- down regulated, green – metabolites of interest. Data was analyzed using the online MetaboAnalyst tool, after log transformation and Pareto scaling of normalized positive and negative-mode combined datasets.

Figure S1. **Assessment of differential metabolic pathways between embryonic and maternal livers at 3h post polyI:C by untargeted metabolomics.**

A Pathway analysis of significantly different metabolites (up/down) between embryonic (eLiver) and maternal liver (mLiver) for data presented in Fig 6C as assessed by untargeted metabolomics at 3h post polyI:C injection and induction of MIA. Level 1-3 confidence metabolites were used for this analysis. For Level 3 compounds, metabolite annotation corresponding to the first hit to an online database (Metabolika or ChemSpider) was used. Data was analyzed using the online MetaboAnalyst tool, after log transformation and pareto scaling of normalized positive and negative-mode combined datasets.
